# Psychedelic enhancement of flexible learning weeks after a single dose

**DOI:** 10.1101/2024.12.17.629035

**Authors:** Elizabeth J. Brouns, Tyler G. Ekins, Omar J. Ahmed

## Abstract

Psychedelic drugs have shown therapeutic potential for the treatment of multiple neuropsychiatric disorders chiefly by promoting long-lasting plasticity in the prefrontal cortex (PFC). A critical function of the PFC is the ability to apply previously learned rules to novel scenarios, a skill known as cognitive flexibility. Here, we show that a single dose of 25CN-NBOH – a serotonin 2A receptor-preferring psychedelic – improves performance on a relatively complex flexible reversal learning task in mice, measured 2-3 weeks after the dose. This effect was seen in both male and female mice. This behavioral finding complements previous cellular results showing that a single psychedelic dose induces long-term structural changes in the PFC and uniquely demonstrates sustained improvements in cognitive flexibility in a novel behavioral paradigm weeks after the initial psychedelic dose in mice. This high throughput task also provides a rapid, automated way to assess other candidate psychedelics for their impact on cognitive flexibility in mice.

## INTRODUCTION

Psychedelic drugs have been successfully used to treat multiple neuropsychiatric disorders, including major depressive disorder, post-traumatic stress disorder, and substance use disorders (1–14). These neuropsychiatric disorders are precipitated by chronic stress, which leads to both structural and functional changes in the prefrontal cortex (PFC) in humans and rodents (15–22). The therapeutic potential of psychedelics may be due to the ability to restore neural circuits damaged in these pathologies by boosting synaptic activity (23–31).

The PFC is responsible for controlling many cognitive functions, including working memory, memory retrieval, decision-making, and executive function (32,33). One key aspect of executive function is the ability to apply previously learned rules to novel situations, which is also known as cognitive flexibility (33,34). Flexibility disruptions are associated with neuropsychiatric disorders, such as depression and post-traumatic stress disorder (PTSD), as well as neurodevelopmental and neurodegenerative disorders (34,35). Cognitive flexibility has been examined using tasks such as the Flanker Task, Stroop Task, and the Wisconsin Card Sorting Task, however these kinds of tasks are largely limited to humans (36). In contrast, most cognitive flexibility tasks for rodents involve attentional set-shifting paradigms, which involve the learning of two separate rules and associated cues, and reversal learning, which involves applying a learned rule to a reversed scenario (37,38). While attentional set-shifting paradigms are popular with rats, reversal learning is a simple yet effective method for cognitive flexibility in rodents, including mice (38–41). Reversal learning paradigms can be extremely diverse, from the kind of task being taught to the number of reversals involved in the paradigm, and these details are critical when evaluating previous literature’s examination of cognitive flexibility through reversal learning.

As single psychedelic administrations promote structural changes in the PFC that last for several weeks(24,29,31), here we asked if a single psychedelic administration could also induce a weeks-long enhancement of flexible learning ability in mice. We conducted a reversal learning task in which female and male mice were administered a single dose of a psychedelic drug or saline and found enhanced performance on the reversal task which persists for at least 3 weeks after one psychedelic dose.

## METHODS

### Animals & Behavioral Apparatus

The open source, programmable Feeding Experimentation Device version 3 (FED3) device was used for all behavioral experiments. We programmed the FED3 via Arduino to deliver a 10mg reward pellet if an animal successfully pressed the correct sequence of nose poke holes (e.g. left-right or right-left within 30 seconds, depending on the phase of the task). The reward pellets used in this study were 10 mg Bio-Serv Dustless Precision Pellets in the chocolate flavor. The cages used with the FED3 devices were modified standard mouse housing cages, with holes drilled into the front of the cage and magnets affixed to the cage’s front to allow the animal to interact with the reward well and nose poke holes and ensure the device stays flush to the side of the cage during data collection. Data were collected within the animals’ vivarium on a static shelf to minimize any effect of changing locations and ensure ample room for both the cage and the device on the shelf. The animals’ vivarium ran on a reversed light cycle with lights off (dark phase) from 7:00 AM to 7:00 PM. Each day, data were collected from approximately 10:00 AM to 2:00 PM, within the vivarium’s dark phase. After each 4-hour session, the animals were returned to their home cages until the next day. 27 adult male and female C57BL/6 mice with a mean age of ∼6 months but the data from one mouse were excluded because the mouse did not interact with the FED3 device. All procedures were approved by the University of Michigan Committee on the Use and Care of Animals.

### Procedure

Animals were injected intraperitoneally with either 25CN-NBOH (N=12 mice) a 5-HT_2A_ receptor-preferring agonist, at a dose of 10 mg/kg, or saline to function as a control (N=14 mice). To allow the blinding of the main experimenter, these injections were done by another experimenter, and the main experimenter remained blinded until after the protocol had been completed. After injection, the animals were left to rest for 24 hours in their home cage before beginning an 85% free feeding weight schedule for 2 days. During those two days, a few reward pellets were dropped into each animal’s cage. This was done to ensure proper food motivation and acclimation to the pellets.

After 2 days of food restriction, animals began a 5-day training period. For 2 days the animals underwent the habituation phase of the protocol, in which the animals were introduced to the FED3 and experimentation cages. Over the course of two separate 4-hour sessions, the FED3 automatically delivered a reward pellet every 4 minutes and in response to any pokes to either nose poke hole. After the two days of habituation, the animals began fixed-ratio 1 (FR1) training phase in which a reward pellet would be delivered any time the mouse poked the left nose poke hole. Similarly, these sessions were 4 hours long each and took place over the course of 3 days.

After the five days of training were completed, the mouse was then introduced to the sequential fixed-ratio 2 (SEQFR2)-forward and SEQFR2-reversal phases. To receive a reward pellet in the SEQFR2-forward phase, the animal must poke the left nose poke hole followed by the right nose poke hole within 30 seconds. Should the animal poke the right hole in isolation, the left hole twice in a row, or not follow a left poke with a right poke, the device entered a 10 second timeout phase in which no further pokes would be registered and any nose pokes during this timeout period were ignored. Like all other sessions, these sessions lasted for 4 hours each over the course of 6 days. After those 6 days elapsed, the SEQFR2 rule was reversed, meaning the animal had to poke the right nose poke hole first followed by the left to receive a reward pellet. Like the forward phase, the reversal phase lasted for 6 days. After the protocol was completed, mice were returned to their normal feeding schedules, cages were cleaned and sanitized with Liquinox lab detergent, and FED3s were sanitized with 70% ethanol. The experimental timeline and protocol overviews are summarized in Figure 1.

**Figure 1.**
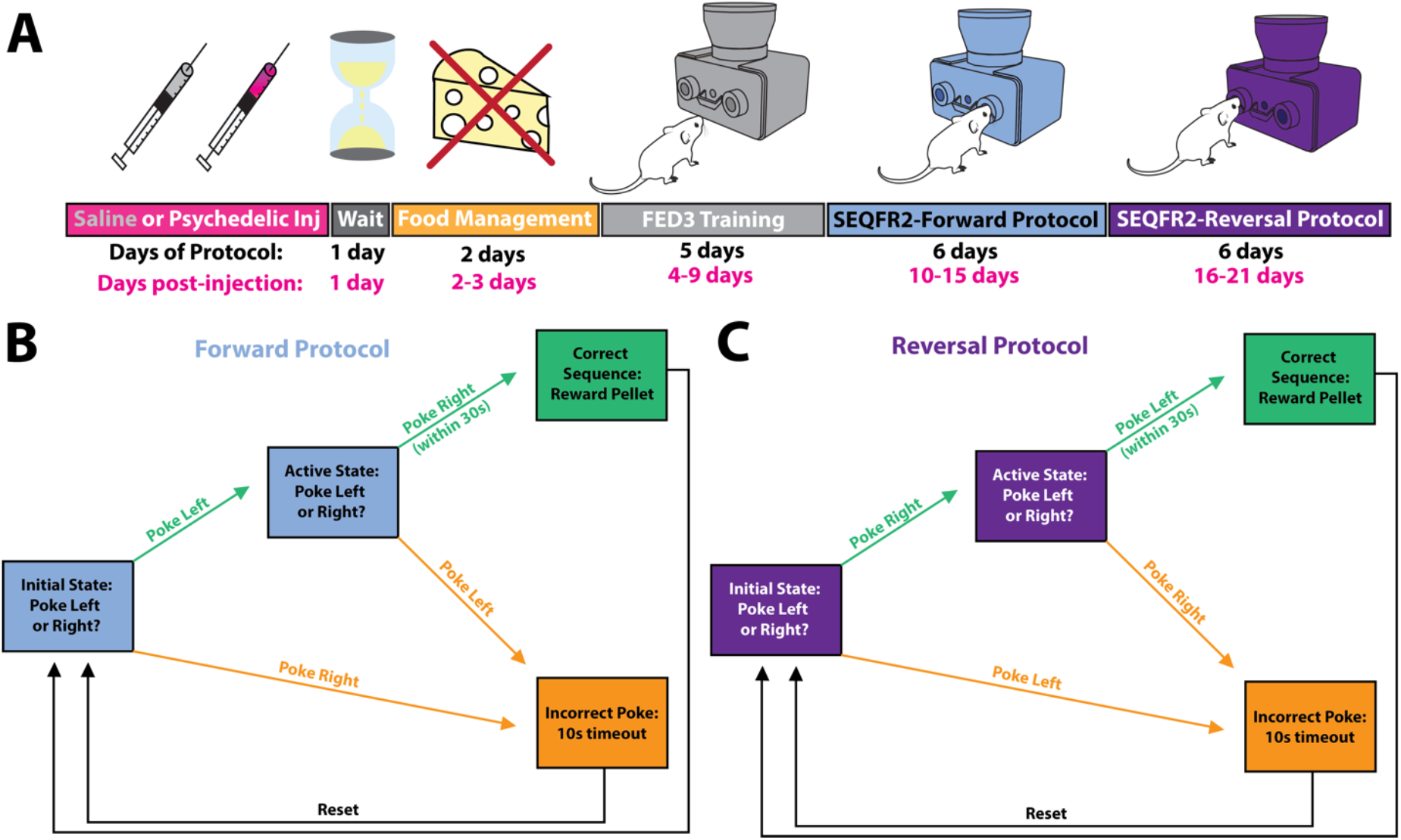
Experimental timeline and overview. (**A**) Experimental timeline^61^. (**B**) Schematic of the SEQFR2-forward protocol. Mice poke left and then poke right within 30 seconds to earn a reward pellet. (**C**) Schematic of the SEQFR2-reversal protocol. Mice now are required to poke right then left within 30 seconds to get a reward pellet.

### Behavioral Analysis

In the SEQFR2 task, poke efficiency is the main measure of performance. This was calculated by finding the ratio of pellets dispensed out of all registered pokes done by the animal in each hour. As an additional metric of behavioral performance, we examined cumulative pellets dispensed over the course of each phase, as well as the proportion of correct trials initiated out of all trials (“percent correct”). Together, these three metrics reflect the absolute performance animals over the course of the task (cumulative pellets), as well performance relative to the amount of engagement with the device (poke efficiency and percent correct).

### Statistics

Statistical procedures were performed with Prism GraphPad (version 10.3.0). Statistical test information and significance are provided in each figure legend.

## RESULTS

To determine if psychedelic treatment induces long-lasting changes in flexible learning ability, we treated female and male mice with a single dose of the serotonin 2A (5-HT_2A_) receptor-preferring agonist 25CN-NBOH (42–45) or saline via intraperitoneal (IP) injection. Following a waiting period of one day, light food restriction for 2 days, and 5 days of training with the Feeding Experimentation Device 3 (FED3) device, we utilized a forward sequence learning protocol (Figure 1A). Mice learned to initiate a trial with a left poke and then had to poke right within the subsequent 30 seconds to receive a food pellet (Figure 1B). Following 6 days of 4-hours/day forward protocol sessions, the required sequential poking pattern was reversed. For another 6 days of 4-hour sessions, mice were then required to poke right and then poke left within 30 seconds to receive the food pellet (Figure 1C). This reversal of the experimental protocol is indicative of flexible learning: we measured the degree to which a mouse is able to adapt the previously learned 1 poke/hole sequence rule to a novel situation, which, in this case, was the reversed direction.

We found that psychedelic and saline-treated mice learned the forward task at similar rates, as reflected by the poke efficiency, which represents the proportion of pellets dispensed out of all pokes (Figure 2A), and the percentage of correct trials initiated out of all trials initiated (Figure 2B). While the change in forward learning poke efficiency and percentage correct was not affected by psychedelic treatment (F_(1,557)_=.1721, P=.6784); F_(1,431)_=.7771, P=.3785, the NBOH treated group accumulated more reward pellets than the saline group (F_(1,620)_=7.513, P=0.0063), indicating an increased initiation of trials per hour with the FED3 (Figure 2C), as the baseline and learning rates were similar between groups (Supplemental Figure 1).

**Figure 2.**
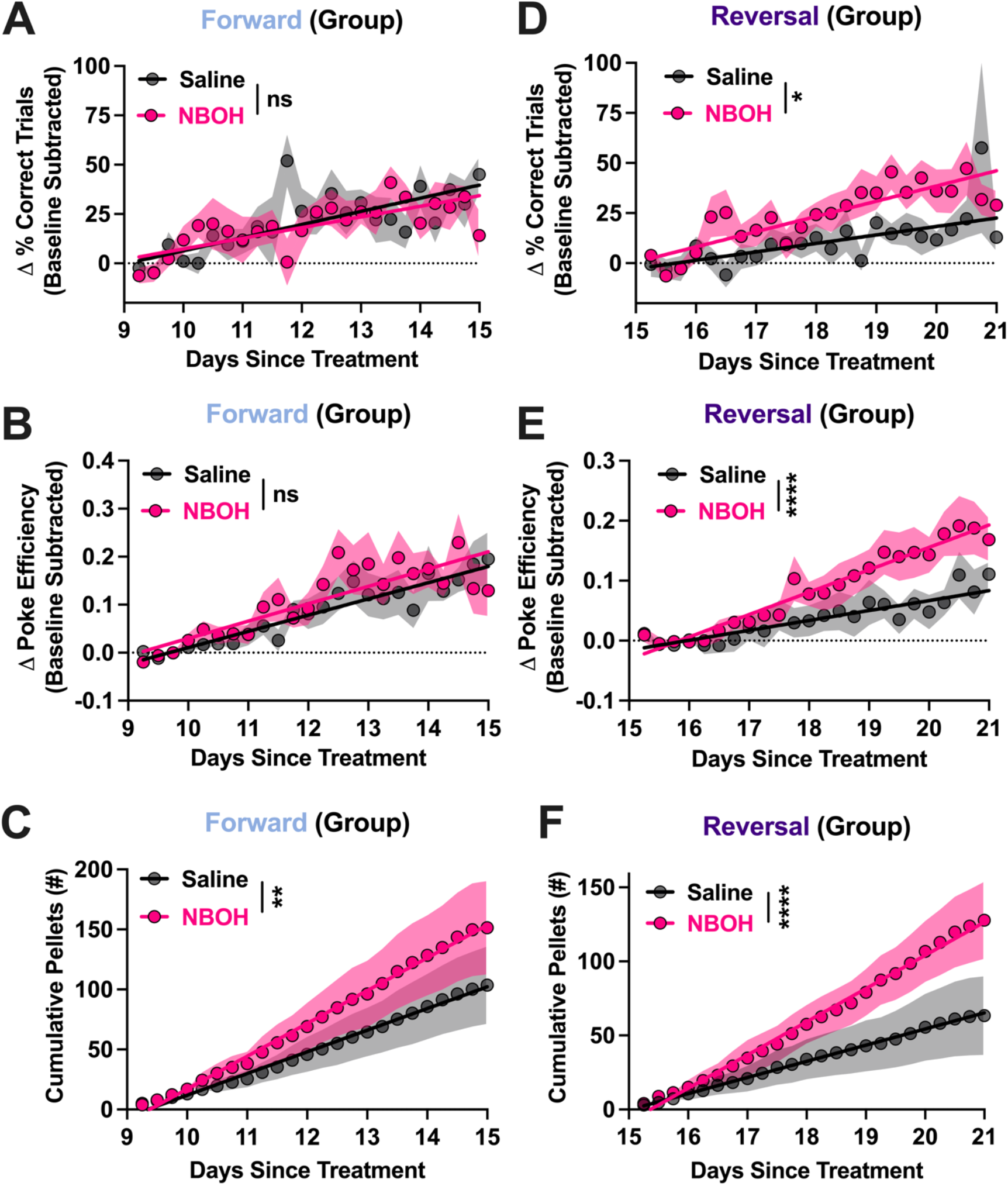
Psychedelic treatment induces a lasting reversal learning ability enhancement. (**A**) Group forward poke efficiency, with no significant effect of NBOH treatment. Each hour-long bin contains 4 points for each hour of the 4-hour per day sessions. (**B**) Group forward percentage of correct trials, indicating no significant effect of NBOH treatment. (**C**) NBOH treatment significantly increased the number of reward pellets dispensed during the forward phase. (**D**) NBOH treatment significantly increased poke efficiency during the reversal phase compared to saline injection, indicating enhanced cognitive flexibility(**E**) NBOH increases the percentage of correct trials initiated. (**F**) NBOH treatment significantly increased the number of reward pellets dispensed during the reversal phase. Shaded regions represent standard error of the mean (SEM), linear regressions shown in pink for NBOH and black for saline; ns, not significant; **p<0.01; ****p<0.0001.

Importantly, during the reversal phase, measured 15-21 days after the single injection, psychedelic treatment resulted in significantly increased learning ability. This is indicated by the increased the efficiency of nose pokes (Figure 2; F_(1,528)_=21.91, P<.0001), the percent correct trials initiated (Figure 2E; F_(1,401)_=6.629, P = .0104), and again by the higher total number of pellets obtained (Figure 2F; F_(1,620)_ =20.74, P<.0001).

Finally, we considered sex as a biological variable to determine if NBOH improved learning in both sexes. We found that, consistent with our sex-independent results (Fig. 2), NBOH treatment did not affect poke efficiency during the forward phase poke efficiency learning during the forward phase (Female: F_(1,334)_=0.986, P=.322; Male: F_(1,219)_=0.004, P=.952), but significantly enhanced poke efficiency during the reversal phase (Figure 3; Female; F_(1,319)_=16, P<.0001; Male: F_(1,205)_=8.3, P=.0044). Thus, psychedelic treatment induced a weeks-long lasting enhancement of reversal learning in both male and female mice.

**Figure 3.**
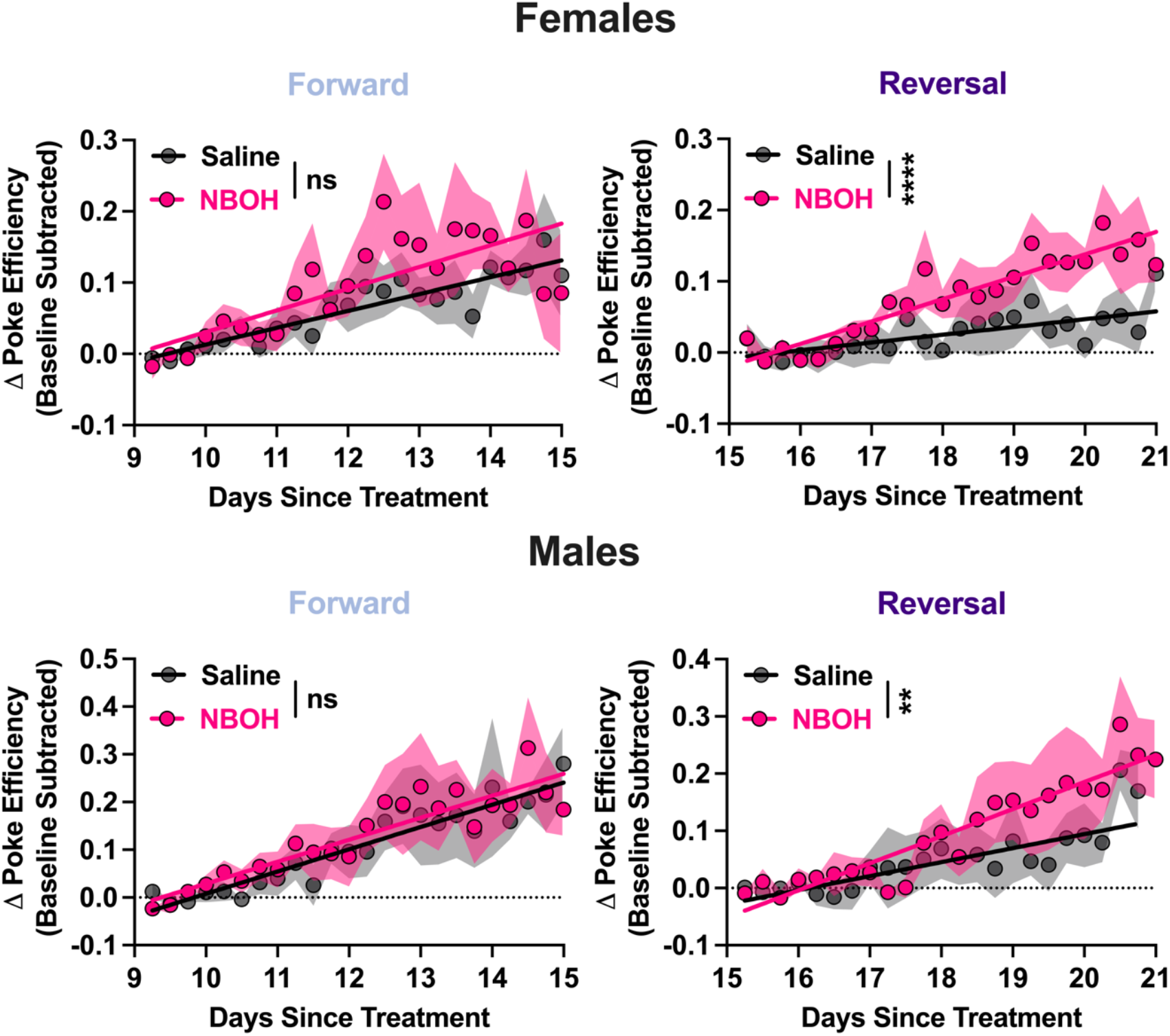
Lasting psychedelic enhancement of reversal learning ability in male and female mice. (**Top**) Forward and reversal changes to poke efficiency indicating NBOH treatment significantly improves reversal learning in female mice weeks after a single dose. (**Bottom**) Forward and reversal changes to poke efficiency in male mice indicating NBOH treatment significantly improves reversal learning weeks after a single dose. Shaded regions represent standard error of the mean (SEM), linear regressions shown in pink for NBOH and black for saline; ns, not significant; **p<0.01; ****p<0.0001.

## DISCUSSION

This study sought to examine the effects of a single psychedelic dose on flexible learning in both female and male mice. We found that even 2-3 weeks after a single dose, NBOH significantly enhanced reversal learning ability. Poke efficiency, percentage of correct trials, and cumulative pellets dispensed were all improved during the reversal phase in mice that received NBOH compared to mice that received saline. No significant effects were observed during the forward phase in terms of poke efficiency or correct trial percentage, but there was a significant increase in the number of pellets obtained, indicating increased drive to engage with the task.

This study contrasts with previous preclinical psychedelic reversal learning studies in terms of drug administration timepoints (46–50). We administered the psychedelic or saline control 15 days before the start of the reversal protocol. Thus, our study focuses on the longer-term therapeutic effects of the psychedelic drug. It is important to distinguish such longer-term effects from immediate or short-lasting acute effects, that may be more related to the mind-altering impact of psychedelics and not to their longer-term therapeutic effects. In a two-choice visual discrimination task, 25CN-NBOH (1-2 mg/kg) was found to have no significant effects on reversal learning in mice when administered acutely, immediately before testing (49). Other previous rodent studies using attentional set-shifting and T-maze paradigms found impairment of flexible learning with acute administration of the psychedelics DOI (1 mg/kg) or 25CN-NBOH (1 mg/kg) on cognitive flexibility (48,50). However, one study found the enhanced cognitive flexibility with acute psilocybin (1 mg/kg) in the same attentional set shifting paradigm that found impairment following administration of DOI (50). The differences in the acute effects of psychedelics on reversal learning may be due to the study design, discussed below, as well as a combination of drug and dose. DOI and 25CN-NBOH are much more potent (>10x) than psilocybin (42,43,51–53). Concentration-dependent acute suppression working memory (27,54–56) likely explains why the relatively less potent psychedelic psilocybin doesn’t acutely impair behavioral performance at this dose. As the long-term effects of psychedelics on cognition are the effects that are most relevant therapeutically, it is important that future work continues to examine the sustained, in addition to acute, effects of psychedelics.

The current study also contrasts with previous studies in protocol design. We selected a reversal learning paradigm as they are sufficiently complicated and somewhat easier to interpret compared to attentional set-shifting or T-maze paradigms. Two recent studies have also made use of reversal learning paradigms and have found long-term (study 1: 3 days; study 2: 14 days) positive effects of psychedelics on cognitive flexibility in female rats, but there are a few notable differences in behavioral protocols than the current study (46,47). As opposed to conducting a sequential FR2-style task with only one reversal of the task, the 14-day long study implemented an FR1 task that reversed every 10 successful trials, and while they demonstrate that psilocybin increases the number of successes over time, they do not show if psilocybin improves accuracy, or if this is a function of increased trial initiations after psilocybin (46). Our FR2 style task with a single reversal after many days of training highlights the different effects psychedelics have on initial (forward) learning and reversal learning separately, which we would have otherwise been unable to do in a paradigm that frequently reverses. Our paradigm also likely results in fewer random successes that could inflate an animal’s actual performance as we require precisely two pokes in two separate holes. In addition, we conducted the task in both female and male mice. A similar study done testing the long-term effects of DOI on reversal learning found that, depending on task structure, DOI has mixed effects on reversal learning ability (57). A week-long evaluation of initial learning after dosing appeared to assist in the enhancement of flexible learning, while if the animals were not further exposed to the task prior to reversal after dosing, DOI appears to have a negative effect on reversal learning ability. This finding suggests that further work needs to be done to evaluate what role further practice following dosing has on cognitive flexibility.

In humans, psilocybin treatment has been found to improve cognitive flexibility up to one month after dosing (13,14). However, these studies utilized a within-subject repeated measures design with no non-psilocybin control group (13), or with low dose psilocybin (1 mg) as the control group (14). Although promising, it is possible behavioral performance was improved through familiarity with the task design rather than a direct result of the psychedelic treatment. It is currently unknown if a single psychedelic dose would improve cognitive flexibility measured in a human study using independent measures.

PFC neurons have been shown to undergo spinogenesis and synaptogenesis after a single psychedelic administration through a pathway requiring 5-HT_2A_ receptor activation (23–31). Here, we use 25CN-NBOH, which is 50-100x more selective for 5-HT_2A_ receptors over 5-HT_2B_ and 5-HT_2C_ receptors and has even weaker affinity for other 5-HT receptors (42,43). Future research into the long-term effects of other psychedelic drugs on cognitive flexibility will need to be conducted to examine if psychedelics that target additional 5-HT receptor subtypes have similar long-lasting effects, or to determine if the interaction with other 5-HT receptors abolishes the ability to enhance long-lasting flexibility. In addition, it remains unknown if non-hallucinogenic 5-HT_2A_ receptor agonists such as 2-bromo-LSD, lisuride, and 6-fluoro-diethyltryptamine (58– 60), are also able to induce a lasting enhancement of flexible learning.

Psychedelic-mediated weeks-long enhancement of reversal learning ability allows for many further directions of research. Future studies could examine the effects psychedelics may have on mice across different ages. Additional studies could also determine the effects of different psychedelic drugs, dose levels, number of doses, or dose timing in this behavioral paradigm. While we did find an enhancement in a mouse model of cognitive flexibility with a 5-HT_2A_ receptor-preferring agonist, we did not use 5-HT_2A_ knockout mice to see if the absence of 5-HT_2A_ receptors would cause any deficits in this behavior, or if psychedelics would improve this behavior without the engagement of 5-HT_2A_ receptors. The task design presented here will facilitate future studies that can address these and other questions. This will allow for an even greater mechanistic understanding of the relationship between psychedelic treatment and cognitive flexibility.

## ACKNOWLEDGEMENTS

This work was supported by: NIH R01MH129282; NIH P50NS123067, NIH T32DA007268, University of Michigan Eisenberg Family Depression Center Eisenberg Scholar Award.

## SUPPLEMENTARY FIGURE

**Supplemental Figure 1.**
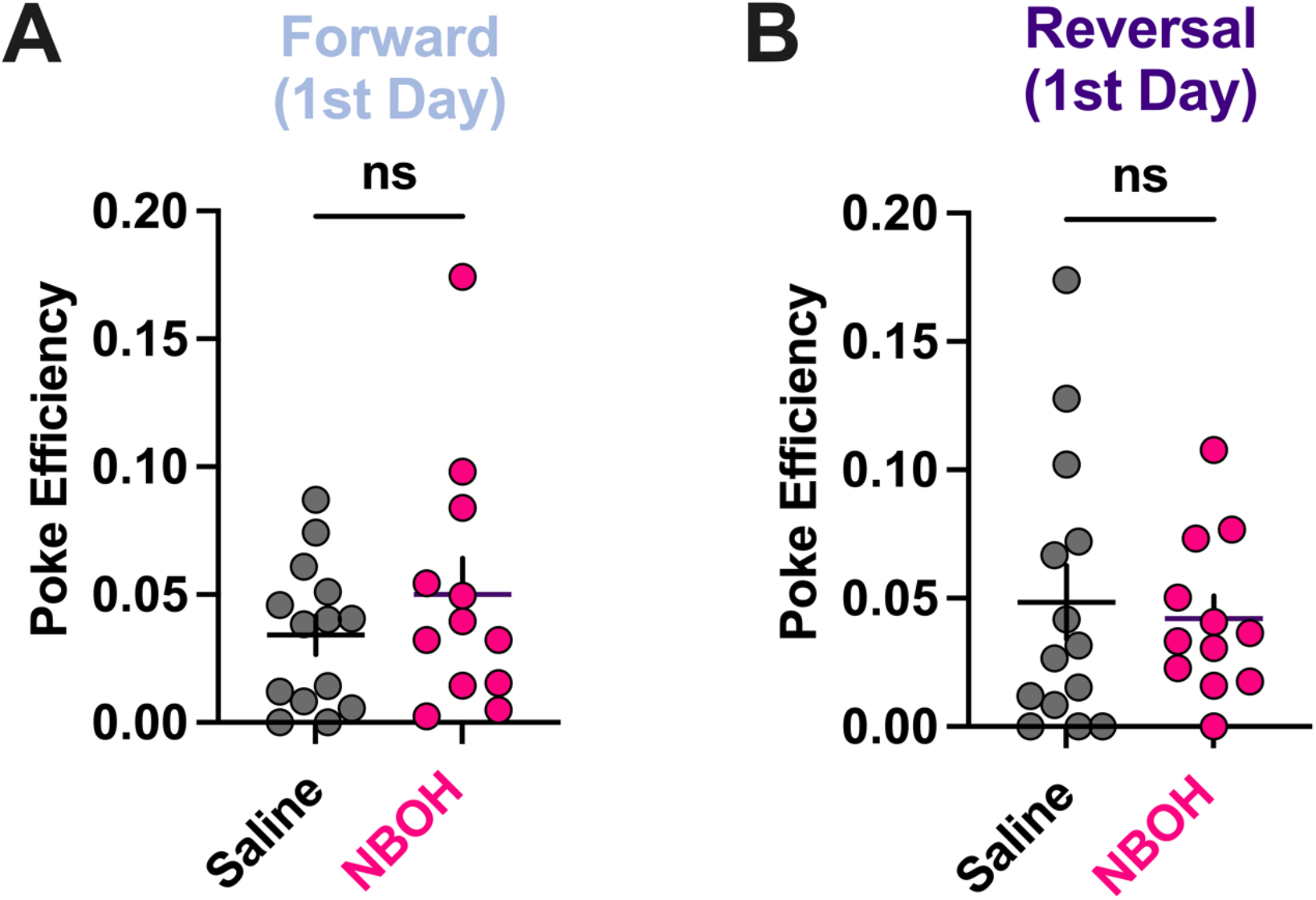
Starting forward and reversal poke efficiency is similar between treatment groups. **(A)** Forward starting (Day 9) poke efficiency is not different between saline and NBOH groups. **(B)** Reversal starting (Day 15) poke efficiency is not different between saline and NBOH groups. Bars and error bars represent mean and standard error of the mean (SEM). Statistics indicate unpaired two-tailed t-test between groups, ns, not significant. Forward: t_(24)_=1.034, P=.3312; Reverse: : t_(24)_=0.3595, P=.7223.

